# A mixed nitrogen diet and compartmentalized utilization for *Mycobacterium tuberculosis* replicating in host cells: results of a systems-based analysis

**DOI:** 10.1101/542480

**Authors:** Khushboo Borah, Martin Beyß, Axel Theorell, Huihai Wu, Piyali Basu, Tom A. Mendum, Katharina Nöh, Dany JV Beste, Johnjoe McFadden

**Affiliations:** Faculty of Health and Medical Sciences, University of Surrey, Guildford, GU2 7XH, UK; School of Veterinary Medicine, University of Surrey, Guildford, GU2 7XH, UK; Forschungszentrum Jülich GmbH, Institute of Bio- and Geosciences, IBG-1: Biotechnology, 52425 Jülich, Germany

**Keywords:** *Mycobacterium tuberculosis*, host macrophages, intracellular pathogen, nitrogen metabolism, systems biology, isotopic labelling

## Abstract

Nitrogen metabolism of *Mycobacterium tuberculosis* (Mtb) is crucial for the survival and virulence of this pathogen inside host macrophages but little is known about the nitrogen sources acquired from the host or their route of assimilation. Here we performed a systems-based analysis of nitrogen metabolism in intracellular Mtb and developed ^15^N-Flux Spectral Ratio Analysis (FSRA) to probe the metabolic cross-talk between the host cell and Mtb. We demonstrate that intracellular Mtb acquires nitrogen from multiple amino acids in the macrophage including glutamate, glutamine, aspartate, alanine, glycine and valine, with glutamine being the predominant nitrogen donor. Each nitrogen source is uniquely assimilated into specific intracellular pools indicating compartmentalised metabolism. This was not observed for *in vitro*-grown Mtb indicating that there is a switch in nitrogen metabolism when the pathogen enters the intracellular environment. These results provide clues about the potential metabolic targets for development of innovative anti-tuberculosis therapies.

## Introduction

Tuberculosis (TB) is one of the top 10 causes of morbidity and mortality in the human population responsible for 1.6 million death and 10 million new infections every year (World health organization, 2018; Flynn, 2006; Zumla et al., 2013). The causative agent of TB, Mtb is primarily resident within the hostile environment provided by the phagosome of macrophages and can therefore resist stress conditions such as low pH, hypoxia, reactive oxygen species, reactive nitrogen species and nutrient starvation (Gouzy et al., 2014a; Russell, 2001; Rustad et al., 2009). Metabolism within this restricted niche is key to the survival and pathogenesis of Mtb (Beste et al., 2011; Ehrt and Rhee, 2013; Eoh et al., 2017; Warner, 2015). Understanding Mtb metabolism within the macrophage has therefore become an important focus for research, with the aim of identifying vulnerable metabolic pathways which could be targeted with drugs. Carbon metabolism in Mtb has been intensively investigated and host derived lipids, cholesterol, and CO_2_ have been identified as essential nutrients for intracellular growth and survival in animal models of TB (Beste et al., 2011; Beste et al., 2013; Ehrt et al., 2018; Eisenreich et al., 2010; Gouzy et al., 2014b; McKinney et al., 2000; Niederweis, 2008; Schnappinger et al., 2003). Our ^13^C-Flux Spectral Analysis (FSA) based approach applied to the intracellular carbon metabolism of Mtb also demonstrated that several nonessential amino acids are imported into the phagosome highlighting their availability as potential nitrogen sources within the macrophage. However, the identity of the primary nitrogen sources for intracellular Mtb remains uncertain. Amino acid acquisition and metabolism is important for the pathogenesis of several intracellular bacterial pathogens, including Mtb (Das et al., 2010; Eylert et al., 2008; Gouzy et al., 2013; Gouzy et al., 2014c; Kloosterman and Kuipers, 2011). For example intracellular *Salmonella enterica* Serovar Typhimurium and *Streptococcus pneumoniae*, requires arginine for intracellular survival and virulence (Das et al., 2010; Kloosterman and Kuipers, 2011), demonstrating that this amino acid is the nitrogen source. *Listeria monocytogenes* has been shown to acquire aspartate, alanine and glutamate from the host macrophages as the source of nitrogen for its *de novo* amino acid synthesis (Eylert et al., 2008). *In vitro*, Mtb can utilize the amino acids alanine, arginine, asparagine, aspartate, glutamate, glutamine, glycine, isoleucine, proline and serine and also urea as sole nitrogen sources (Lin et al., 2012; Lofthouse et al., 2013). However, there is very little information of what nitrogen sources are available to intracellular Mtb growing within a macrophage.

The Mtb genome encodes several transporters for both inorganic and organic nitrogen compounds; *e.g*. ammonium chloride (Amt), nitrate (NarK2); as well as ABC type amino acid transporters (Cole et al., 1998). Aspartate and asparagine have been detected in the Mtb phagosome and using mutagenesis approaches, the researchers showed that the transport and assimilation of aspartate was not required for intracellular survival but was essential for survival in a murine model of TB whereas asparagine had an essential role in acid stress resistance in the host cell (Gouzy et al., 2013; Gouzy et al., 2014a). An Mtb mutant strain deficient in the aspartate transporter AnsP1 also had defects in the synthesis of nitrogen containing compounds indicating that aspartate is a potential nitrogen donor (Gouzy et al., 2014a). Glutamate biosynthesis is essential for mycobacterial growth as glutamate dehydrogenase (GDH), the enzyme that catalyses production of glutamate from 2-oxoglutarate, is essential for the intracellular survival of Mtb (Cowley et al., 2004; Gallant et al., 2016; Viljoen et al., 2013); however GDH is also required for resistance to acidic and nitrosative stress in *Mycobacterium bovis* BCG so it is unclear what the role of glutamate is during intracellular survival. Glutamine synthetase (GS), the enzyme that synthesizes glutamine from glutamate is also essential for intracellular growth and virulence of Mtb (Tullius et al., 2003), suggesting that insufficient supplies of this amino acid are available from the intracellular environment. Mtb auxotrophic mutants of leucine, proline, tryptophan, and glutamine have previously been shown to be severely attenuated *in vivo* (Hondalus et al., 2000; Lee et al., 2006; Smith et al., 2001), indicating that biosynthesis of these amino acids is required in the intracellular environment; however several other auxotrophic mutants (e.g., methionine, isoleucine or valine) can successfully proliferate in macrophages, indicating that these amino acids can be acquired from the macrophage milieu (Awasthy et al., 2009; McAdam et al., 1995). However, none of these studies have directly measured uptake and assimilation of nitrogen by intracellular Mtb.

Isotopic tracer studies, combined with direct interpretation of the labelling patterns emerging in intracellular metabolites, are a powerful means to derive information on nutrient contributions to the production of different metabolites (Buescher et al., 2015). We previously used ^13^C isotopomer analysis, and developed ^13^C-FSA to identify the intracellular carbon sources (Beste et al., 2013). However this approach was unsuitable for the current study because of the lack of backbone rearrangements in the metabolic network as compared to carbon, and limited mass isotopomer information from single nitrogen atoms. To overcome these limitations, here we developed ^15^N-FSRA, a safeguarding method to counteract “over-ambitious” reasoning, that accounts for all possibilities that are in-line with the labelling data and biochemistry including reaction reversibilities in an otherwise unbiased manner. This computational platform allowed for an unbiased analysis of all potential amino acid uptake systems. Specifically, to counteract the limited measurement information, an array of ^15^N labelling experiments was conducted under equal conditions which was simultaneously evaluated in an integrative system-based approach. We demonstrate that intracellular Mtb utilises multiple nitrogen sources and that nitrogen assimilation is compartmentalized and distinct from *in vitro* growth. These results improves our understanding about the nitrogen assimilation of this important pathogen whilst also providing tools which can be applied to other intracellular pathogens.

## Results

### Mtb co-assimilates multiple nitrogen sources during intracellular growth

To measure uptake and assimilation of nitrogen during the intracellular growth of Mtb, THP-1 macrophages were infected with Mtb in the presence of a ^15^N tracer-aspartate (Asp), glutamate (Glu), glutamine (Gln), leucine (Leu), alanine (Ala) or glycine (Gly) for 48 h. Asparagine was not tested as it had previously been shown to be a nitrogen source for intracellular Mtb (Gouzy et al., 2014a). Using an identical method to that developed by Beste et al., 2013, macrophage and Mtb fractions were separated, and ^15^N incorporation into protein-derived amino acids from infected macrophages and intracellular Mtb was measured using GC-MS. Amino acid pairs-glutamate, glutamine and aspartate, asparagine cannot be distinguished using GC-MS, so, the ^15^N enrichments for these amino acid pairs are combined together as glutamate/glutamine (Glu/n) and aspartate/asparagine (Asp/n) respectively. Experimental controls of uninfected THP-1 macrophages and *in vitro* grown Mtb were cultivated in RPMI with each of the tracers for 48 h.

These experiments established that the ^15^N labelling profile of protein-derived amino acids from intracellular Mtb was distinct from that of Mtb grown in RPMI (Figure S1, Figure 1C). For example the labelled nitrogen from ^15^N_1_-Leu was incorporated predominantly into Leu when Mtb was grown in RPMI, but was widely dispersed into other amino acids for Mtb growing intracellularly (Figure S1D). Similarly, for RPMI grown Mtb, the label from ^15^N_1_-Asp, ^15^N_1_-Glu and ^15^N_2_-Gln was incorporated into Asp/n and Glu/n. However during intracellular growth, labelled nitrogen from ^15^N_1_-Asp was incorporated into almost all measured amino acids including Asp/n and Glu/n (Figures S1A-S1C). The distinct ^15^N labelling profiles of intracellular vs RPMI-grown Mtb demonstrated that there was negligible cross contamination from extracellular Mtb in these experiments. Moreover there was no cross-contamination during the separation of the macrophage and the intracellular Mtb as evidenced by the unique labelling profile of protein-derived amino acids from these two fractions (Figures 1A and 1C). The labelling profiles of protein-derived amino acids from infected and uninfected macrophages were virtually identical indicating that Mtb infection does not significantly perturb the nitrogen metabolism of macrophages (Figures 1A and 1B).

**Figure 1.**
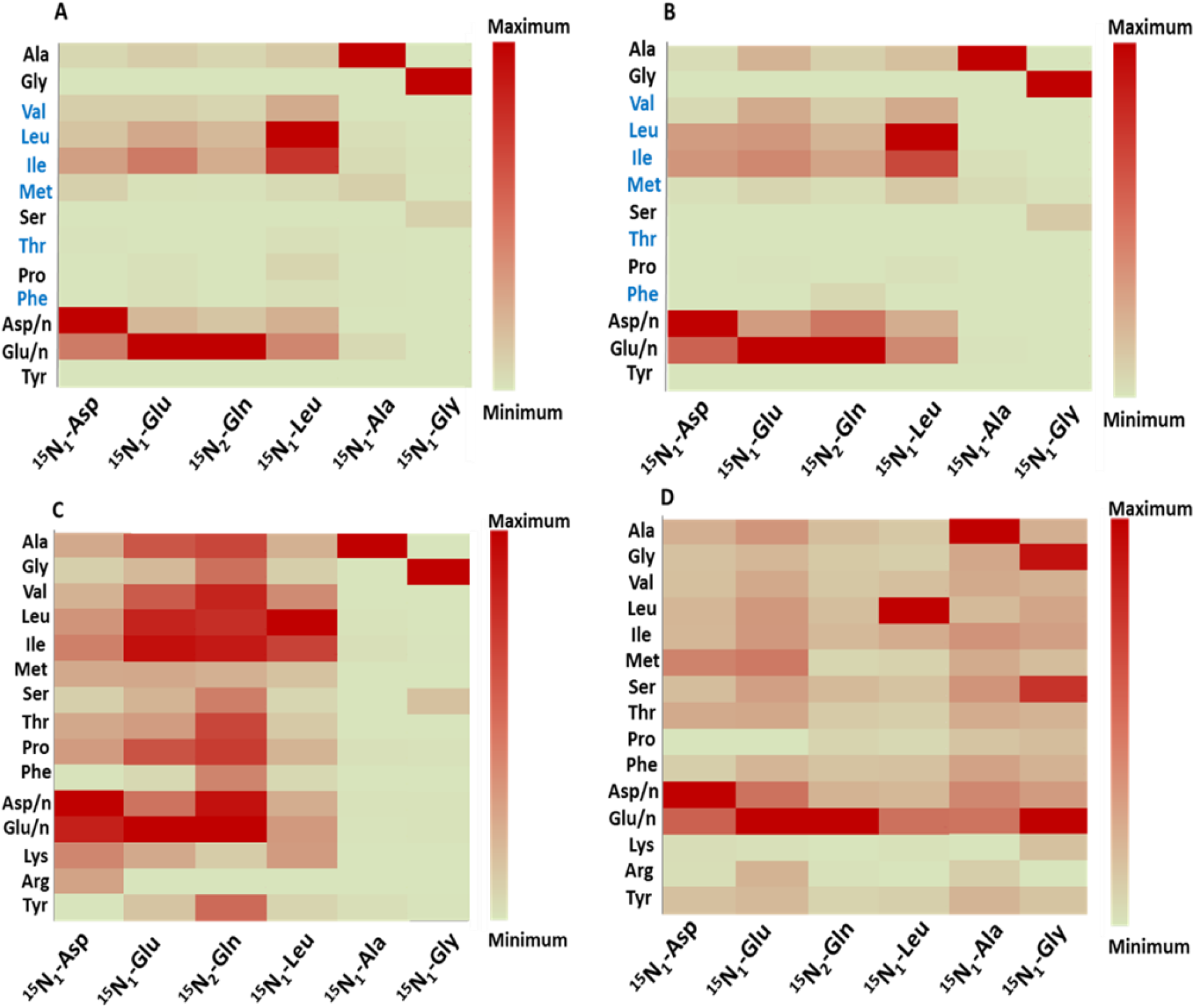
Assimilation pattern of different nitrogen sources. Heat maps are shown for amino acids derived from Mtb-infected macrophages (A), uninfected macrophages (B), intracellular Mtb (C) and *in vitro*-grown Mtb in rosins minimal media (D). Assimilation pattern are shown for 6 tracers-^15^N_1_-Asp, ^15^N_1_-Glu, ^15^N_2_-Gln, ^15^N_1_-Leu, ^15^N_1_-Ala and ^15^N_1_ Gly. Essential amino acids for macrophages are highlighted in blue. Heat maps were produced using enrichment (%) of amino acids recorded for each of the individual tracer experiments in data file S1. The maximum and minimum enrichments for each heat map are described using colour keys. Abbreviations for amino acids alanine (Ala), glycine (Gly), valine (Val), leucine (Leu), isoleucine (Ile), methionine (Met), serine (Ser), threonine (Thr), proline (Pro), Phenylalanine (Phe), aspartate/asparagine (Asp/n), glutamate/glutamine (Glu/n), lysine (Lys), arginine (Arg) and tyrosine (Tyr).

In accordance with expectations, most of the amino acids derived from macrophages remained unlabelled reflecting their direct uptake from unlabelled nitrogen sources in the tissue culture media rather than from any transamination from the tracer (Figure 1A). This is not the case for intracellular Mtb. For example labelled nitrogen from either ^15^N_1_-Asp, ^15^N_1_-Glu or ^15^N_2_-Gln was incorporated predominantly into Asp/n, Glu/n and isoleucine in the macrophage but labelling was measured in nearly all amino acids derived from intracellular Mtb (Figures 1A and 1C). This data also reveals potential directions of transamination in Mtb. For example, nitrogen from ^15^N_2_-Gln was assimilated into six amino acids, Asp/n, Glu/n, isoleucine (Ile), Leu, valine (Val) or Ala in infected macrophages (Figure 1A); whereas in Mtb, 8 additional amino acids were labelled (Figure 1C). As the label from Gln has already been transaminated into several amino acids in the macrophage, any of those amino acids could be the ultimate source of the Mtb’s labelled nitrogen.

### Nitrogen metabolism is compartmentalised in Mtb

Our results also demonstrate that nitrogen was assimilated differently depending on the original amino acid source in intracellular Mtb. We use the term compartmentalisation to refer to the selective transfer of nitrogen from the tracers to other amino acids (Figure 1C). For example, nitrogen from ^15^N_1_-Asp, ^15^N_1_-Glu, ^15^N_2_-Gln and ^15^N_1_-Leu was incorporated into many additional amino acids, but nitrogen from ^15^N_1_-Ala and ^15^N_1_-Gly was incorporated predominantly into only Ala, Gly and serine (Ser) pools respectively. The enrichment profiles for amino acids obtained from the tracers were also distinct, for example, the maximum ^15^N from ^15^N_1_-Asp was transferred to glutamate; ^15^N_1_-Glu to Ile; ^15^N_2_-Gln to Asp/n and ^15^N_1_-Leu to Ile. Also, some amino acids were preferentially labelled by particular tracers. For example, labelling in tyrosine (Tyr) indicates that the nitrogen is transferred from ^15^N_1_-Glu/^15^N_2_-Gln but not from ^15^N_1_-Asp or any other tracers used. Similarly, nitrogen in arginine (Arg) can be donated by ^15^N_1_-Asp but not from any other tested amino acid.

To investigate whether the compartmentalized nitrogen distribution is restricted to intracellular growth of Mtb, we performed similar ^15^N tracer experiments with Mtb grown in Roisins minimal media with glycerol and ammonium chloride as the carbon and nitrogen source respectively, with the addition of ^15^N labelled tracers. As can be seen, the results confirmed the compartmentalized distribution of ^15^N from each amino acid, even during *in vitro* growth (Figure 1D, Figure S2). Also noteworthy is the promiscuous pattern of distribution of nitrogen from both 15N_1_-Ala and ^15^N_1_-Gly *in vitro*, which is very different from the restricted distribution found for intracellular Mtb (Figures S3E-S3F). This result could reflect the scarcity of these amino acids within the phagosome niche.

Out of all the nitrogen sources tested, nitrogen from ^15^N_2_-Gln was most widely distributed to other amino acids in intracellular Mtb, suggesting that Gln is the principle nitrogen donor for Mtb during intracellular growth (Figure 1C). The comparison of enrichment profiles between 15N_1_–Glu and ^15^N_2_-Gln demonstrated that both tracers gave similar ^15^N distribution pattern in amino acids derived from uninfected macrophage and infected macrophage (Figures 1A and 1B); but ^15^N_2_-Gln provided significantly higher enrichments than^15^N_1_-Glu for the majority of the intracellular Mtb amino acids (Figure 1C). Direct comparison of ^15^N_1_–Glu and ^15^N_2_-Gln in Figure 2 demonstrated that there were no significant differences in enrichments between ^15^N_1_– Glu and ^15^N_2_-Gln in the macrophage (Figure 2A). But in intracellular Mtb, ^15^N_2_-Gln gave significantly higher levels of ^15^N enrichments for majority of the amino acids. For example, Ala, Gly, Val, Leu, Ile, Ser, threonine (Thr), proline (Pro), phenylalanine (Phe) and Asp/n were all highly labelled with ^15^N_2_-Gln compared to ^15^N_1_-Glu.

**Figure 2.**
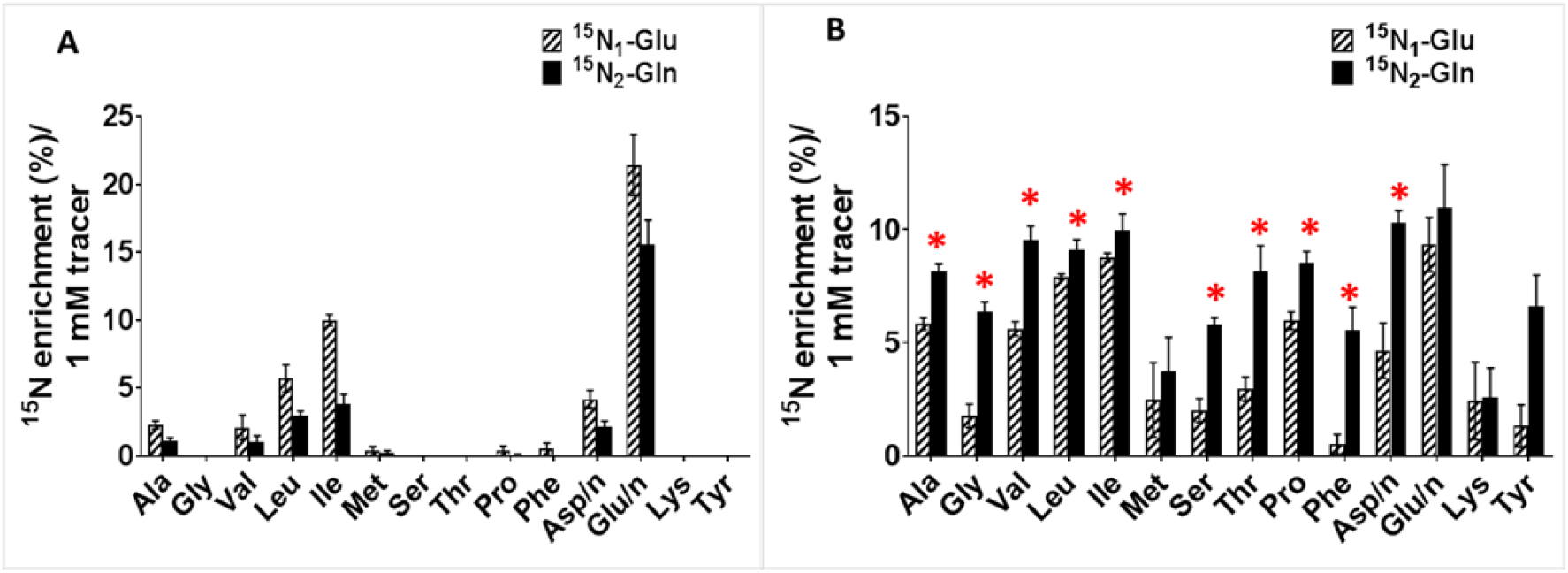
Comparison of ^15^N enrichment from the tracers-^15^N_1_-Glu and ^15^N_2_-Gln. Data is shown for infected macrophages (A) and intracellular Mtb (B). Enrichments are shown for the amino acids-Ala, m/z 260; Gly, m/z 246; Val, m/z 288; Leu, m/z 274; Ile, m/z 274; Met, m/z 320; Ser, m/z 390; Thr, m/z 404; Pro, m/z 258; Phe, m/z 336; Asp/n, m/z 418; Glu/n, m/z 432; Lys, m/z 431; Tyr, m/z 466 measured using GC-MS. Values are mean ± SEM from 3-4 biological replicates. Statistically significant differences were calculated using Holm-Sidak multiple t-tests; **\***, p < 0.001.

### Uptake vs *de novo* synthesis in intracellular Mtb

To further evaluate the role of each macrophage amino acid pool in the provision of nitrogen to intracellular Mtb and to normalize for different levels of label in each of the macrophage pools, we calculated the Nitrogen Incorporation Ratio (NIR) between ^15^N enrichment into intracellular Mtb amino acids compared to the same amino acid in the macrophage. Amino acids acquired directly from the host cell that are not subject to any additional synthesis or metabolism will tend to have similar ^15^N enrichment as the macrophage amino acids. *De no*vo synthesis or additional metabolic processes, such as transamination, in Mtb will be revealed by different levels of ^15^N enrichment in Mtb compared to the macrophage. For example, if, as well as being imported, an Mtb amino is also *de novo* synthesized by Mtb, then the amount of ^15^N label in Mtb’s amino acid pool will depend on the level of label of the nitrogen source for that *de novo* synthesis. If the source has more ^15^N label than the imported amino acid then the Mtb pool will be enriched for ^15^N relative to the macrophage pool. If the source has less ^15^N than the imported amino acid then the Mtb pool will be depleted for ^15^N relative to the macrophage pool. The analysis thereby provides a direct, data-driven measure of whether amino acids are net donors or recipients of nitrogen.

Figures 3A-3G shows the ratio of ^15^N enrichment for 7 amino acids derived from intracellular Mtb as compared with amino acids derived from infected macrophages. A NIR of 1 for amino acids such as Ala, Val and Ser indicates that these amino acid are acquired directly from the macrophage and then directly incorporated into Mtb’s biomass (Figures 3E-3G). However, when ^15^N-Asp was used as label for the Asp/n pool, the NIR was less than 1 (Figure 3A) indicating that any Mtb Asp/n imported from the macrophage was being diluted with nitrogen from a source that was less labelled than macrophage Asp/n pool. However, when ^15^N_2_-Gln was used as label, the ratio for the Asp/n pools was > 4 indicating that any Mtb Asp/n imported from the macrophage was being diluted with nitrogen from a source that was at least 4 x more labelled than macrophage Asp/n. The same was not true when ^15^N_1_-Glu was the source of label. The obvious conclusion is that the Asp’s nitrogen in Mtb is largely derived from macrophage Gln. The NIR for Leu was < and/or > 1 from ^15^N_1_-Leu and other tracers, suggesting that Leu was not directly taken up from the macrophage and that nitrogen in Leu was *de novo* synthesized from other nitrogen sources (Figure 3C). The NIR for Val was 1 from 15N_1_-Leu tracer, suggesting that nitrogen from Leu is transaminated to Val that is directly acquired from the macrophage by Mtb (Figure 3E). For the majority of amino acids including Asp/n, Leu, Ile, Val and Ala, the ratio was also significantly higher (up to 8 fold) when ^15^N_2_-Gln was used as the source of label. The ratios were not determined for Gly and Ser when heterologous tracers were used, because there was no enrichment for these amino acids in macrophage. But the labels for these amino acids in Mtb were highest, when ^15^N_2_-Gln was used (see Figure 1C), thus confirming Gln to be the predominant nitrogen donor for intracellular Mtb for the tested amino acids.

**Figure 3.**
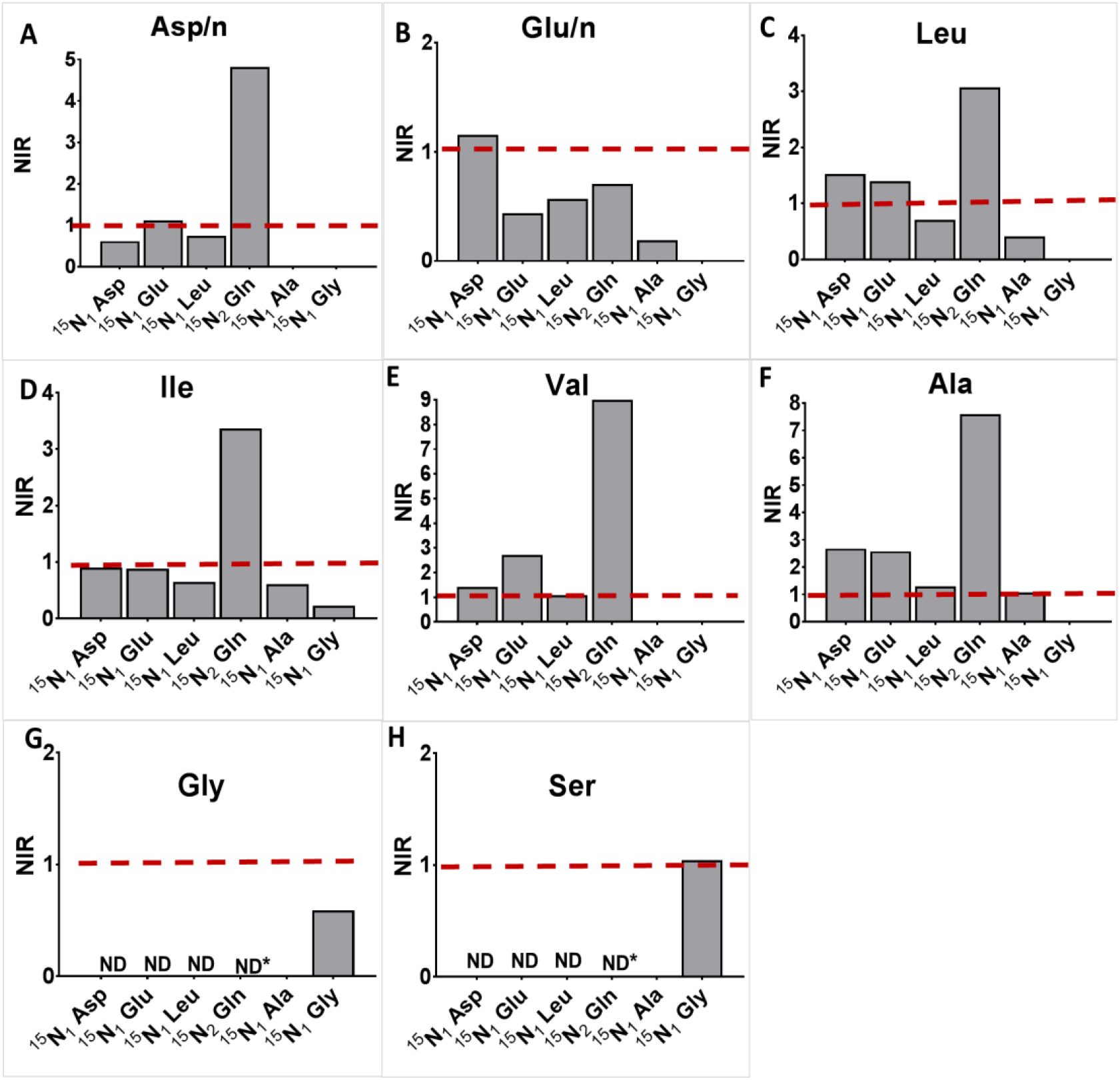
Intracellular Mtb uptakes nitrogen sources from macrophages. The NIR of ^15^N enrichment (%) in intracellular Mtb to infected macrophages are shown for Asp/n, m/z 418 (A), Glu/n, m/z 432 (B), Leu, m/z 274 (C), Ile, m/z 274 (D), Val, m/z 288 (E), Ala, m/z 260 (F), Gly, m/z 246 (G), Ser, m/z 390 (H). A NIR of 1 indicates that the particular amino acid in Mtb was acquired directly from the macrophage). NIRs were calculated from the enrichments measured from 3-8 individual macrophage infection experiments using the tracers-^15^N_1_-Asp, ^15^N_1_-Glu, ^15^N_2_-Gln, ^15^N_1_-Leu, ^15^N_1_-Ala and ^15^N_1_-Gly. Ratios that were not determined for Gly (G) and Ser (H) are indicated by ND and the maximum ^15^N detected in Mtb from ^15^N_2_-Gln is indicated by *.

Comparison of Figure 1 and Figure 3 also reveal additional interesting features. For example, it is apparent from Figure 1 that nitrogen from ^15^N_1_-Ala and ^15^N_1_-Gly is not promiscuous, as the label from ^15^N_1_-Ala remained with Ala and from ^15^N_1_-Gly remained with Gly and Ser pools from both the macrophage and Mtb. Neither is therefore a nitrogen donor for either the macrophage or Mtb. However, it is also apparent that *de novo* synthesis is also contributing to the Mtb’s Ala pools, since ^15^N from other labelled amino acids is able to reach the Mtb’s Ala pool. Gly however shows a very different pattern as homologous labelling with ^15^N_1_-Gly gave a NIR less than 1 indicating that the Mtb’s Gly pool is being diluted by a less-labelled nitrogen source but not any of the tested amino acids since none of them gave significant levels of incorporation into Gly. The NIR of Ser was 1 for label from Gly, consistent with a synthesis pathway in which imported Gly is converted to Ser, possibly by glycine hydroxymethyltransferase.

### Development of ^15^N-FSRA to provide a possibilistic view on the direct uptake of nitrogen sources by Mtb

Although qualitative conclusions such as above, can be drawn from simple labelling patterns generated by ^15^N_1_-Ala and ^15^N_1_-Gly (Figures 4(I) and 4(II)), it is limited by the complexity of the data. For the labelling patterns generated by ^15^N_2_-Gln, there are multiple possible solutions for uptake patterns, for example, whether the ^15^N in Mtb’s Glu/n is incorporated directly from the macrophage pool of ^15^N-Glu/n or indirectly via another labelled macrophage amino acid and then transaminated to Glu/n (Figure 4(III). The NIR does not account for the intracellular redistribution of nitrogen. But there is more in the mass spectrometry data than the aggregated enrichment information; and there is known biochemistry of nitrogen metabolism. The idea is to bring together mass isotopomer patterns with a network model that describes the flow of label on the level of atoms. Here we neither can use traditional ^13^C MFA since we want to infer the uptake nor our former ^13^C-FSA tool since we have to take too many possible uptakes to account. We therefore developed ^15^N-FSRA to model the nitrogen metabolism of intracellular Mtb. This approach inputs data from the ^15^N tracer experiments into an *in silico* model of central nitrogen metabolism in Mtb, but without making prior assumption on the number or combination of potential nitrogen sources. The method calculates the range of potential explanations in terms of nitrogen uptake flux relative to biomass contribution which is consistent with the available labelling data sets simultaneously. We call this range flux spectral range (FSR). The FSR of an amino acid gives, thus, an unbiased possibilistic measure reflecting the possible nitrogen flows.

**Figure 4.**
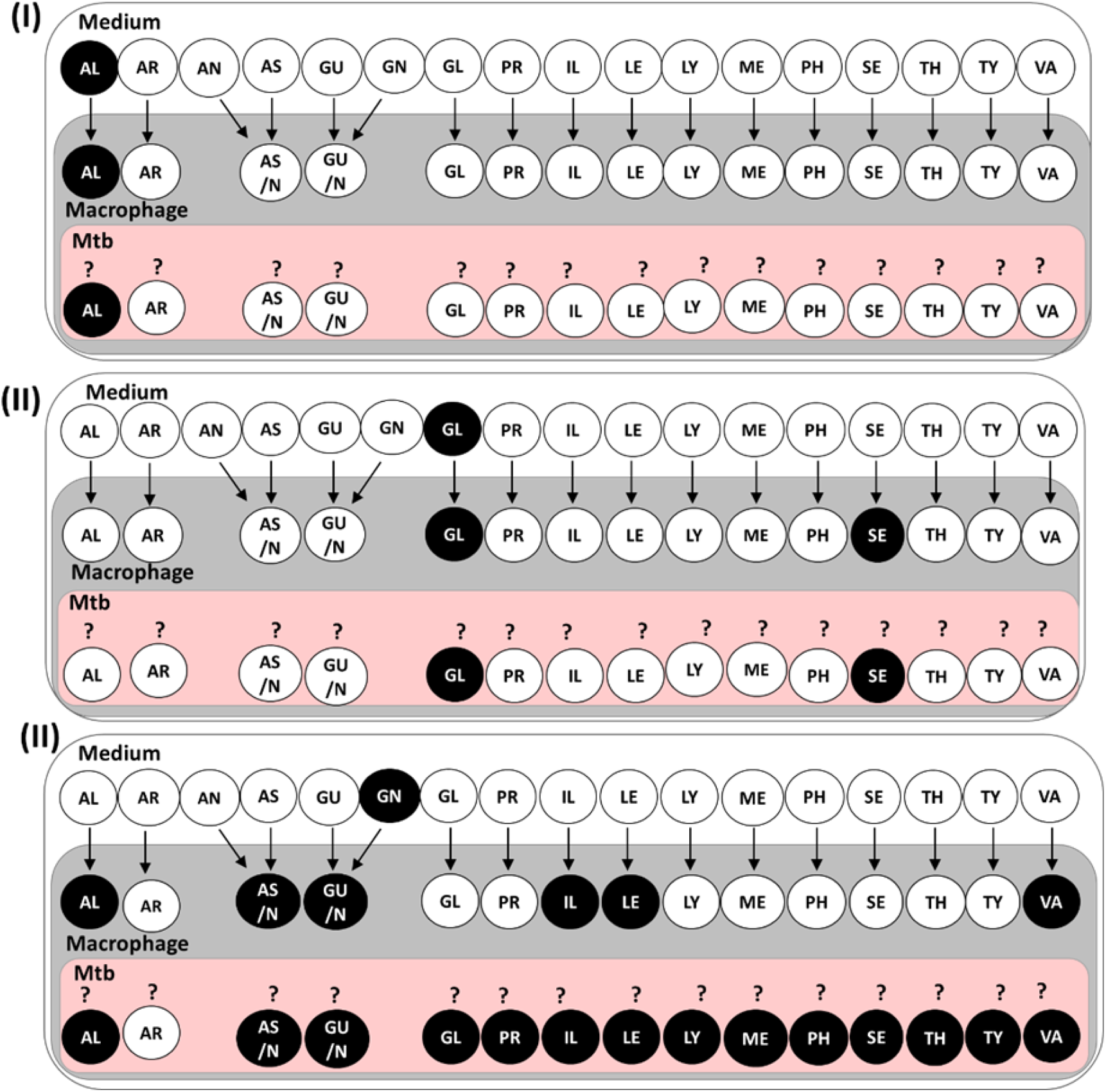
Qualitative ^15^N Labelling patterns of intracellular Mtb, infected macrophages and growth medium. The pattern is shown for the tracers-^15^N_1_-Ala (I), ^15^N_1_-Gly (II) and ^15^N_2_-Gln (III). Labelled amino acids are filled in black and the unlabelled amino acids in white. Amino acids acquired directly from the medium by infected macrophage are indicated by arrows. The uncertainty of direct or indirect uptake of amino acids by Mtb from the macrophage is indicated by ?. Abbreviations for amino acids-AL (alanine), AR (arginine), AN (asparagine), AS (aspartate), AS/N (aspartate/asparagine), GU (glutamate), GN (glutamine), GU/N (glutamate/glutamine), GL (glycine), PR (proline), IL (isoleucine), Le (leucine), LY (lysine), ME (methionine), PH (phenylalanine), SE (serine), TH (threonine), TY (tyrosine) and VA (valine).

To this end, a ^15^N metabolic model was constructed for Mtb including nitrogen atom transitions and reaction reversibilities (supplementary data file S2, supplementary Figure S4). The model was able to analyse the amino acids Glu/Gln and Asp/Asn as separate pools, respectively and was solely constrained by a biomass equation taken from our previous work (Beste et al., 2005; Beste et al., 2011). The uptake of ^15^N labelled amino acids by Mtb (in total 20) was modelled by deconvolution of the macrophage derived amino acid mass isotopomer distributions. The measurements of all six tracer experiments were incorporated into the model. From the labelling data measurements, Histidine (His) and Arg were excluded due to low detection of these amino acids by GC-MS. After global flux fitting, the relative FSRs (minimum ratio, median, and maximum ratio) of amino acid nitrogen uptake flux to the nitrogen content of each amino acid in biomass were deduced (supplementary data file S3). For this analysis amino acids with minimum/maximum FSR∼1 indicated that this amino acid is available in the macrophage and fully incorporated into Mtb’s biomass whereas a minimum/maximum FSR of 0 indicates that the amino acid is unavailable and must be completely synthesised *de novo* by Mtb. A minimum/maximum FSR > 1 indicates that the uptake of the amino acid is greater than the biomass requirements, and therefore that this amino acid is a likely nitrogen donor. It should be noted that amino acids with FSRs between 0 and 1 may still be nitrogen donors, but the deficit in their uptake compared to biomass requirement must be made up by *de novo* synthesis with an alternative nitrogen source.

Using this analysis 14 amino acids (minimum ratio > 0) are available and taken up by intracellular Mtb from the macrophage (Table 1, supplementary Figure S6). Cysteine, Arg, His and tryptophan FSRs are not included in the analysis due to the lack of experimental measurements for these amino acids. 9 amino acids had FSRs between 0 and 1, and the biomass nitrogen requirement was met by *de novo* synthesis. The narrow FSR around 1 for Ala confirmed our previous analysis that this amino acid was taken up directly by Mtb from the macrophage and incorporated directly into biomass. Furthermore, the analysis showed that Gln and Val are nitrogen donors for the synthesis of other amino acids with high certainty (min FSR > 1) while the computational results for Glu, Gly, and Ser (min FSR < 1 and max FSR > 1) remained inconclusive. For Asp and Gly the minimum FSR was > 0 and maximum > 1, suggesting that these amino acids were both acquired and synthesized *de novo*, but may also be a nitrogen donor. The minimum FSR was 0 for Glu and Ser, indicating no uptake from the macrophage. The FSR median of Glu is close to 0 (Table 1, supplementary data file S4), implying that it is unlikely that Glu is taken up by substantial amounts and that was not a nitrogen donor for intracellular Mtb. The median for Ser was > 1, suggesting that Ser could be nitrogen donor, with the nitrogen being transaminated from another nitrogen source. Furthermore, our results revealed that 9 amino acids (Asn, Ile, Leu, Met, Lys, Tyr, Phe, Pro, and Thr) are synthetized *de novo* to suffice biomass requirements for growth in varying proportions, as seen from the narrow ratio ranges.

**Table 1.** Flux spectral ranges determined with ^15^N FSRA. Minimum and maximum FSR for all computationally accessible amino acids, and median of their distributions (Supplementary data file S3). Amino acids are grouped into five categories: “↓” – spectral ranges in the interval [0,1) indicate that the biomass nitrogen need has to be fulfilled by *de novo* synthesis of the amino acid; “?” – spectral range is essentially 1, nitrogen biomass requirement is balanced with uptake, no nitrogen is shared; “0” – the amino acid is not be taken up; “?” – spectral range is inconclusive, the amino acid might be synthesized *de novo*, but may also be a nitrogen donor; “?” – spectral ranges are larger then 1, indicating that this amino acid is available as a nitrogen donor.

|  | Nitrogen source | Flux spectral range |  |  |
| --- | --- | --- | --- | --- |
|  |  | Minimum ratio | Maximum ratio | Median |
| ↓ | Asn | 0.06 | 0.19 | 0.13 |
| ↓ | Pro | 0.31 | 0.38 | 0.34 |
| ↓ | Thr | 0.37 | 0.49 | 0.43 |
| ↓ | Ile | 0.52 | 0.64 | 0.55 |
| ↓ | Tyr | 0.57 | 0.63 | 0.60 |
| ↓ | Leu | 0.58 | 0.96 | 0.61 |
| ↓ | Met | 0.64 | 0.77 | 0.68 |
| ↓ | Phe | 0.63 | 0.75 | 0.69 |
| ↓ | Lys | 0.78 | 0.82 | 0.81 |
| → | Ala | 0.98 | 1 | 1 |
| 0/○ | Glu | 0 | 1.63 | 0.02 |
| 0/○ | Ser | 0 | 3.59 | 1.31 |
| ○ | Gly | 0.51 | 3.17 | 0.67 |
| ○ | Asp | 0.61 | 12.46 | 2.11 |
| ↑ | Gln | 1.50 | 11.85 | 2.47 |
| ↑ | Val | 1.82 | 13.45 | 2.88 |

## Discussion

We and others have shown that Mtb co-metabolises multiple carbon sources during intracellular growth in the human host cell (Beste et al., 2013; Ehrt et al., 2018; McKinney et al., 2000). Here, we demonstrate that this is also the case for nitrogen. Using ^15^N isotopomer profiling and the newly-developed ^15^N-FSRA systems-based inference tool, we describe, for the first time, the nitrogen metabolic phenotype of Mtb inside human macrophages (Figure 5). With the new systems-based ^15^N-FSRA tool, we predicted that Mtb has access to most amino acids within the macrophage (Table 1) and this uptake was insufficient to meet biomass requirement for most tested amino acids and therefore must be complemented with *de novo* synthesis from nitrogen donors. This analysis identified the amino acids-Asp, Glu, Gln, Val, Ala and Gly as the intracellular nitrogen sources utilised by Mtb in human macrophages.

**Figure 5.**
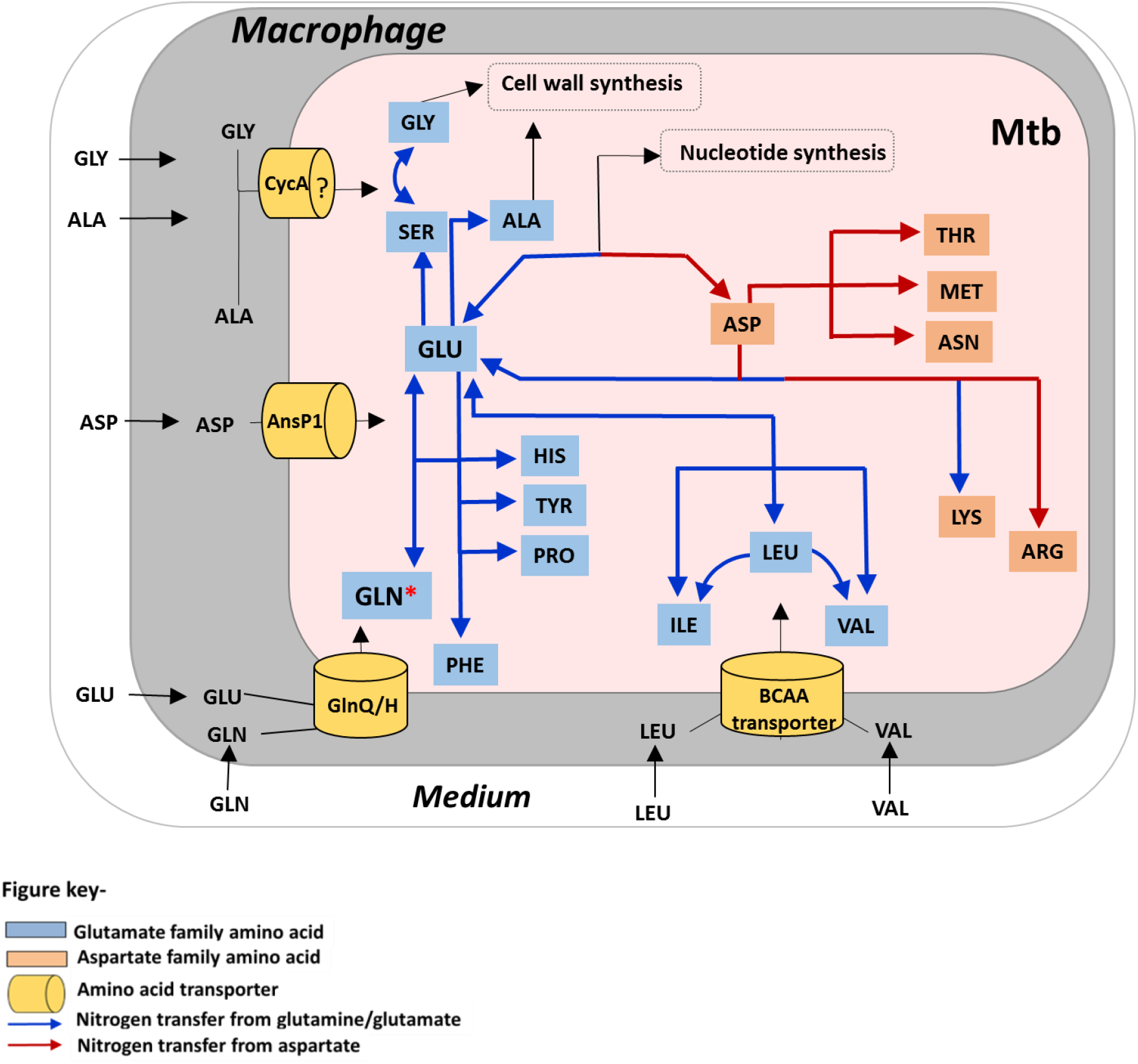
Schematic representation of nitrogen metabolism (acquisition and assimilation) in intracellular Mtb. Macrophages acquires nitrogen sources-ASP (aspartate), GLU (glutamate), GLN (glutamine), LEU (leucine), ALA (alanine) and GLY (glycine) directly from the growth media. GLU/GLN is taken from the host via yet unidentified transporter. ASP is accessible to intraphagosomal Mtb, which it uptakes from the host macrophages via AnsP1 (Gouzy et al., 2014a). LEU, ILE (isoleucine) and VAL (valine) are acquired from the host macrophages via yet unindentified branched chain amino acid probably ABC-type transporter. ALA and GLY are possibly acquired via CycA transport system. GLN, VAL and ASP were potential nitrogen donors for cellular protein synthesis with GLN as the principle nitrogen donor in intracellular Mtb and is indicated by **\***. Nitrogen from GLN was transaminated primarily to GLU and ASP for the synthesis of other amino acids-ALA, GLY, SER (serine), HIS (histidine), TYR (tyrosine), proline (PRO), phenylalalnine (PHE), THR (threonine), MET (methionine), ASN (asparagine), LYS (lysine) and ARG (Arginine). ALA and GLY are assimilated mainly into ALA, GLY and SER pools respectively. Limited transamination of nitrogen from ALA and GLY to other amino acids, suggests direct assimilation of these two amino acids for cell wall synthesis.

Further analysis of the data demonstrated that like carbon metabolism in Mtb, (de Carvalho et al., 2010), nitrogen metabolism is similarly compartmentalized but in the case of nitrogen, this is a specific response to intracellular growth. Metabolism has been shown to be compartmentalised in other bacteria with *microcompartments*, such as carboxysomes and metabolosomes, serving to compartmentalize enzymes involved in particular pathways; and to restrict exposure of toxic intermediates to the rest of the cell (Frank et al., 2013). For example, in *Salmonella enterica*, activity of the metabolosome was required to alleviate acetaldehyde toxicity during ethanolamine catabolism (Brinsmade et al., 2005). In the case of intracellular Mtb, different nitrogen sources might be assimilated using enzymes that are similarly localised. For example, enzymes glutamine synthetase and asparaginase are demonstrated to function extracellularly for catabolism of Gln and Asn respectively in Mtb (Harth *et al.*, 1994; Gouzy *et al.*, 2014a) as a strategy to resist acidic stress inside the phagosome.

Earlier studies demonstrated that Asp is an intracellular nitrogen source for Mtb. The pathways for Asp/n assimilation were recently discussed (Gouzy et al., 2013; Gouzy et al., 2014a). However, our results suggest that Gln is the principle source of nitrogen for intracellular Mtb and is the amino acid whose nitrogen is most widely assimilated (Figure 1C). Gln is the major fuel for a variety of mammalian cells including macrophages (Zhang et al., 2017). THP-1 macrophages have previously been shown to assimilate Gln into Glu and were shown to be rich in Glu (Amorim Franco et al., 2017; Zhao et al., 2013; Choi and Park, 2018). In our current and previous study we demonstrated that both Glu and Gln were available to Mtb inside the macrophage (Beste et al., 2013). Here we additionally show that Gln and not Glu is the predominant nitrogen source for intracellular Mtb.

Once in Mtb, Gln can be converted to Glu by glutamate synthase (GltB) (Lee et al., 2018), and the nitrogen transferred to other amino acids by various transaminases (Figure 6). Our results are thereby consistent with previous studies that have demonstrated that Glu/n biosynthesis is essential for intracellular growth and survival of Mtb (Harper et al., 2010; Gallant et al., 2016; Ventura et al., 2013; Tullius et al., 2003). In addition to being the nitrogen donor for other amino acids, Glu/n is also involved in cell wall synthesis and resisting acid and nitrosative stress (Harth and Horwitz, 2003; Wietzerbin-Falszpan et al., 1973; Read et al., 2007). These studies, together with the experiments reported here, establish Glu/n as the hub of nitrogen metabolism in intracellular Mtb. Note that Gln is also involved in modulating host cellular immune responses (Dos Santos et al., 2017), leading to the intriguing hypothesis that Mtb may facilitate its own intracellular survival by metabolising this amino acid. Targeting Glu/n transport and metabolism therefore represents a promising avenue for the development of anti-TB drugs.

Biosynthesis of branched chain amino acids is also essential for the intracellular survival of Mtb (Awasthy et al., 2009). Here we show that the branched chain amino acid Val is acquired directly by intracellular Mtb and is utilised as a nitrogen donor in addition to Gln (Figure 3, Table 1). Val was not directly tested as a tracer for the ^15^N experiments in macrophage and intracellular Mtb, but Val pool was ^15^N labelled from all of the amino acid tracers (except for 15N_1_-Ala and ^15^N_1_-Gly) suggesting that nitrogen was shared between Val and other amino acids. A Val auxotroph of Mtb was able to survive in macrophages (Awasthy et al., 2009); this is consistent with our finding that intracellular Mtb acquired Val from the host. Nitrogen from Val is reversibly transaminated to Glu, Ile and Leu by IlvE, the branched chain transaminase that is required for mycobacterial survival during infection in mice model (Grandoni et al., 1998; Sassetti and Rubin, 2003; Tremblay et al., 2009). The demonstration by Zimmermann et al., 2017 that the biosynthetic genes for branched chain amino acids were downregulated when Mtb was growing intracellularly in macrophages, is consistent with our evidence that these amino acids are also available in the phagosome. Our finding that nitrogen from Leu is assimilated by intracellular Mtb is surprising since a LeuD auxotroph of Mtb was shown to be severely attenuated for growth and virulence in murine bone marrow derived macrophages and in mice, suggesting that Leu was unavailable to Mtb growing within its host (Chen et al., 2012; Hondalus et al., 2000; Sampson et al., 2004). The NIR analysis showed that Leu was not acquired directly from the host but most likely as Val (Figure 3). Also, ^15^N-FSRA showed that Leu uptake is insufficient for Mtb’s biomass requirements, indicating that uptake must be supplemented by *de novo* synthesis, consistent with the auxotrophic results. Note that ^15^N-FSRA also indicated that, to account for the data, there was a minimum Leu uptake, approximately half its biomass requirement, confirming that it is imported as well as *de novo* synthesized.

Our previous work showed that the carbon backbone of Ala was acquired from the host and directly incorporated into Mtb’s biomass (Beste et al., 2013). Here we confirm this result and also show that Ala was not a nitrogen donor for intracellular Mtb (Figure 1C, Table 1). Gly was also acquired from host macrophages but its nitrogen was similarly not widely assimilated. This data suggest that these amino acids could be scarce in phagosome and were used for specific functions. Both Ala and Gly are components of mycobacterial cell wall (Alderwick et al., 2015; Mahapatra et al., 2005; Wietzerbin-Falszpan et al., 1973), therefore, these amino acids acquired from the host were directly incorporated for cell wall synthesis. ^15^N-FSRA also indicated that Ser was not imported during intracellular growth; which is consistent with our earlier observation that is Ser cannot be used as a nitrogen source by Mtb (Lofthouse et. al, 2013). Although this does not mean that import of other amino acids is essential for Mtb intracellular growth, it does suggest that transport systems of these nitrogen sources are potential targets for anti-TB drug development. However, there is only a limited knowledge of the amino acid transport systems in Mtb (Cook et al., 2009). The Asp transporter AnsP1 is essential for the virulence of Mtb in murine models (Gouzy et al., 2013). Although Glu and branched chain amino transport systems are yet to be identified. *Streptococcus mutans* and *Lactococcus lactis glnQHMP* system is known to transport Glu (Krastel et al., 2010; Fulyani et al., 2015), and Mtb encodes GlnQ and GlnP homologues that could function as Glu/n transporters (Cole et al., 1998; Braibant et al., 2000). CycA-the common transporter for Ala, Gly and Ser also remains unstudied (Awasthy et al., 2012; Chen et al., 2012; Cole et al., 1998). Further investigations are required to define these amino transporters of Mtb.

This study is the first description of nitrogen metabolic phenotype of intracellular Mtb, and the development and application of a novel computational systems-based tool, ^15^N-FSRA as a safeguard that can be used to investigate nitrogen uptake and assimilation not only of Mtb but any intracellular pathogen, in complement with direct isotopic labelling interpretation. In conclusion, we have identified that multiple nitrogen sources are acquired and assimilated by Mtb inside the host macrophages, and glutamine is the principle intracellular nitrogen donor. Further investigation of transport and assimilatory pathways associated with these nitrogen sources will provide productive avenues for identification of targets for innovative anti-TB therapies.

## Acknowledgments

This study was supported by the BBSRC grant (BB/L022869/1).

## Author contributions

K.B., D.B., and J.M. conceived and designed the experiments; K.B. performed the experiments and analysed data; M.B., T.A., and K.N. performed FSRA; K.B., D.B., and J.M. wrote the original draft; H.W., P.B., M.B., K.N., and T.A.M contributed to the final draft; D.B., K.N., and J.M. supervised the study.

## Materials and Methods

### Bacterial strains, media and growth conditions

*Mtb* H37Rv and *Mycobacterium bovis* was cultivated on Middle brook7H11 agar and Middlebrook 7H9 broth with 5% (v/v) oleic acid-albumin-dextrose-catalase enrichment supplement (Becton Dickenson) and 0.5% (v/v) glycerol. Roisins minimal media was prepared with the composition described in Khatri et al., 2012 and supplemented with 0.5% glycerol (v/v) as the carbon source and 10mM NH_4_Cl as nitrogen source, 0.1% tyloxapol (v/v) at 37°C with agitation (150 rpm).

### Cultivation of human THP-1 macrophages

The human monocytic cell line was obtained from ATCC TIB-202 and grown in RPMI 1640 medium supplemented with 10% heat inactivated fetal calf serum (sigma) at 37°C, 5% CO_2_ and 95% humidity. For ^15^N labelling experiments, modified RPMI supplemented with 0.2mM L-glutamine was used for testing ^15^N_1_ aspartic acid (sigma), ^15^N_1_ glutamic acid (sigma). RPMI without L-glutamine was used for testing ^15^N_2_ glutamine (sigma). RPMI-1640 with 200 mM L-glutamine was used for testing ^15^N_1_ leucine (sigma), ^15^N_1_ alanine (sigma) and ^15^N_1_ glycine (sigma).

### Infection of macrophages with Mtb

For infection, THP-1 cells and Mtb cultures were prepared as described in Beste et al., 2013. THP-1 cells (3 × 10^7^) were differentiated into macrophages for 3 days using 50 nM Phorbol 12-myristate 13-acetate in a 175 cm^2^ tissue culture flasks. Macrophages were washed with warm PBS supplemented with 0.49 mM Mg^2+^ and 0.68 mM Ca^2+^ (PBS^+^) and 30 ml of RPMI media was added. Mtb was grown in 7H9 broth for a week to an optical density of 1 (1 × 10^8^ colony forming units per ml). Mtb cultures were washed 3 times with PBS and resuspended in RPMI media and added to the macrophages at a multiplicity of infection-5. Each amino acid tracer except for ^15^N_2_ glutamine was then added to the Mtb-infected macrophages at 3 times the concentration of that unlabelled amino acid present in the RPMI. For ^15^N_2_ glutamine experiment, RPMI without unlabelled glutamine was used, because a large amount of unlabelled glutamine was present in RPMI as compared to other amino acids. To reduce the costs of the ^15^N_2_ glutamine labelling experiment, the only glutamine in this RPMI was the tracer itself, added at 1mM concentration which previously demonstrated by Tullius et al., 2003 and also confirmed in this study to have no detrimental effect on the macrophages (data not shown). The final concentration of tracers added during infection were-0.6mM ^15^N_1_ aspartic acid, 0.53mM ^15^N_1_ glutamic acid, 0.8mM ^15^N_1_ leucine, 0.3mM ^15^N_1_ alanine, 1mM ^15^N_2_ glutamine and 0.4mM ^15^N_1_ glycine. Mtb infected macrophages were incubated for 3-4 hours at 37°C, 5% CO_2_ and 95% humidity. After incubation, macrophages were washed 3 times with PBS, tracers were added and left for 48 h at 37°C, 5% CO_2_ and 95% humidity. After incubation for 48 hours, macrophages were washed with cold PBS and Mtb cultures were harvested from macrophages using differential centrifugation (Beste et al., 2013). Amino acid hydrolysates were prepared from macrophage and Mtb using 6M hydrochloric acid and incubation at 100°C for 24 hours. The validity of the method for labelled amino acid extraction from intracellular Mtb was confirmed by comparing the nitrogen assimilation pattern obtained in RPMI grown labelled Mtb (control) (Figure S1). The control was set up by washing Mtb cultures with PBS and resuspending in RPMI for 48 hours with equal amounts of ^15^N isotope tracers that was previously used for infection assays. After 48 hours, Mtb in RPMI were harvested followed preparation of amino acid hydrolysate.

### *In vitro* ^15^N labelling experiments

Mtb was grown in Roisins minimal with media spiked with ^15^N isotope tracers and left for 48 h at 37°C and cultures were agitated at 150 rpm. After 48 hours, Mtb cultures were harvested and amino acid hydrolysate was prepared by boiling Mtb in 6M hydrochloric acid for 24 hours.

### Gas-chromatography mass spectrometry (GC-MS) and ^15^N mass isotopomer analysis

Amino acid hydrolysates were dried and derivatized using pyridine and tert-butyldimethyl silyl chloride (TBDMSCl) (sigma) (Rossi et al., 2017; Masakapalli et al., 2013). Amino acids were analysed using a VF-5ms inert 5% phenyl-methyl column (Agilent Technologies) on a GC-MS system. MS data was extracted using chemstation GC-MS software and were baseline corrected using Metalign (Lommen, 2009). Mass data were corrected for natural isotope effects using Isotopomer network compartmental analysis (INCA) platform (Young, 2014). Average ^15^N in an amino acid was calculated from the fractional abundance of the mass isotopomer in the entire fragment. For this study measurements > 1% were considered to be significantly enriched by the ^15^N tracers.

### Nitrogen network model (of Mtb)

A nitrogen transition model was set up for Mtb using information available in databases (KEGG, BioCyc, Tuberculist) and literature. The network model comprises the amino acid biosynthesis pathways of all 20 protein-derived amino acids, as well as a simplified nucleotide biosynthesis formulation. Reaction reversibilities and requirements for growth were taken from Beste et al., 2005. For 20 protein-derived amino acids unidirectional uptake reactions were formulated while it was assumed that no backflow of nitrogen from Mtb to the phagosome exists. The mass isotopomers observed in the macrophage were deconvolved into distributions of isotopomer species and modelled by additional reactions (Figure S4). For comparability of the inference results across different nitrogen uptake constellations, flux values were formulated relative to the biomass synthesis rate. In total, the model has 98 independent fluxes, from which 50 are intracellular (40 net, 10 exchange) and 48 nuisance fluxes for modelling the deconvolution of the substrate species. This nitrogen transition template model was then duplicated per data set to give rise to a sextuple model (Beyß et al. submitted), where each sub-model shared flux values, biomass requirements and growth rate. Eventually, the sextuple model had 338 free fluxes (50 intracellular and 288 nuisance fluxes) that had to be inferred from 264 independent labeling measurements (15 measurement groups for Mtb and the host, respectively (supplementary file S2, supplementary Figure S4).

### ^15^N Flux Spectral Range Analysis (^15^N-FSRA)

^15^N-FSRA borrows the concept of parallel data integration from COMPLETE-MFA (Leighty et al., 2013) which, however, was reformulated resigning the knowledge of the experimentally inaccessible specific amino acid uptake rates and amending the traditional least squares fitting approach by a tailored regularization approach to punish model complexity. In short, a penalty term was added to the weighted least squares functional

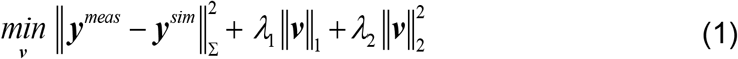

where ***v*** are the (independent) fluxes, ***y***^*meas*^, ***y***^*sim*^ the measured/simulated data, Σ the measurement covariance matrix, and *λ*_*i*_, *i* = 1, 2 regularization parameters punishing non-zero flux values (*λ*_1_ =10.0 and *λ*_2_ = 0.1).

A multi-start optimization strategy was applied to safeguard against local minima while solving Eq. (1). The flux fitting procedure was performed with the high-performance simulator 13CFLUX2 (Weitzel et al. 2013) using the multifitfluxes module with 1,000 randomly sampled starting points and the NAG C optimization library (Version 6.23, Oxford, UK) with a maximum number of 250 iterations. From the 1,000 runs, those with residuals less than a cut-off of 203.5 were accepted (the overall optimal residual value was 182.01, corresponding to a 𝒳^2^ value > 99.99%, supplementary data files S3 and S4). This resulted in 800 accepted fits. From these accepted estimated flux distributions, FSRs (minimum, maximum, median) for each flux were derived (Table 1). Histograms of all FSRs are available in supplementary Figure S5.

